# Tracking Activity In a Deformable Nervous System With Motion Correction and Point-Set Registration

**DOI:** 10.1101/373035

**Authors:** Thibault Lagache, Benjamin Lansdell, Jerry Tang, Rafael Yuste, Adrienne Fairhall

## Abstract

The combination of fluorescent probes with time-lapse microscopy allows for the visualization of the entire neuronal activity of small animals, such as worms or cnidarians, over a long period of time. However, large deformations of the animal combined with the natural intermittency of neuronal activity make robust automated tracking of firing fluorescent neurons challenging. Here we present an hybrid approach where (i) a subset of very bright neurons is used as moving reference points (fiducials) to estimate the elastic deformation of the animal; (ii) deformation is frame-by-frame corrected, and firing neurons are aligned at each time with the initial mask; and (iii) point-set registration is used to robustly track the intermittent activity of all the immobilized neurons. We compare different registration strategies with manual tracking performed over ≈620 neurons over 100 time frames in the cnidarian *Hydra vulgaris*.

**Index Terms:** Fluorescence imaging, wavelet detection, tracking, point-set registration, elastic deformation, Thin Plate Spline (TPS) transform, Coherent Point Drift (CPD), Hydra.

## 1. INTRODUCTION

To understand emergent properties of neuronal ensembles and to decode the neuronal basis of behavior, one would like to record from many, or all, of the neurons in an animal for extended time periods while it is freely behaving [1]. A small set of organisms have been genetically engineered with fluorescent probes and imaged with time-lapse microscopy [2, 3, 4, 5]. The cnidarian *Hydra* is an emerging model system for whole-animal imaging [5], as it is transparent and neurons are sparsely distributed in the body. However, large body deformations and intermittent fluorescence of individual neurons make it challenging to robustly detect and track neuronal activity over a long period of time. Recently, Nguyen et al. developed a hybrid approach to track neurons in the *C. Elegans* worm [6], in which moving neurons are first aligned at each time frame along the principal axis of the stretched animal, and their activity then tracked using point set registration via a Gaussian Mixture Model (GMM) [7]. Here, we adapt this two-step strategy to neuron tracking in *Hydra*, including substantial modifications. Along with large deformations during movement, *Hydra* deforms in two dimensions and cannot easily be transformed into an invariant frame of reference simply using displacement of the principal axis. To our advantage, however, while neurons within the worm are densely packed and imaged in 3D, neurons in Hydra have been imaged in 2D (transparent animal between coverslips) and are relatively sparsely and homogeneously distributed throughout the body. Thus, one should be able to make use of a point-set registration in Hydra that is less computationally intensive than the GMM.

We thus chose to (i) automatically detect firing (fluorescent) neurons at each time frame; (ii) estimate body deformation between two time frames with an elastic transformation, the Thin-Plate Spline (TPS) transform, using a subset of very bright neurons as reference or fiducial points; (iii) iteratively correct for Hydra deformation and align firing neurons with reference frames; and (iv) track the activity of individual neurons with the Coherent Point Drift (CPD) algorithm [8] (Fig. 1). The CPD algorithm ignores spot shape and accounts only for positions. It is thus faster than a GMM and well suited to the distributed (i.e. not densely packed) neural network of Hydra. Finally, we compare different CPD variants against manual tracking on *≈* 620 neurons over 100 frames.

**Fig. 1.**
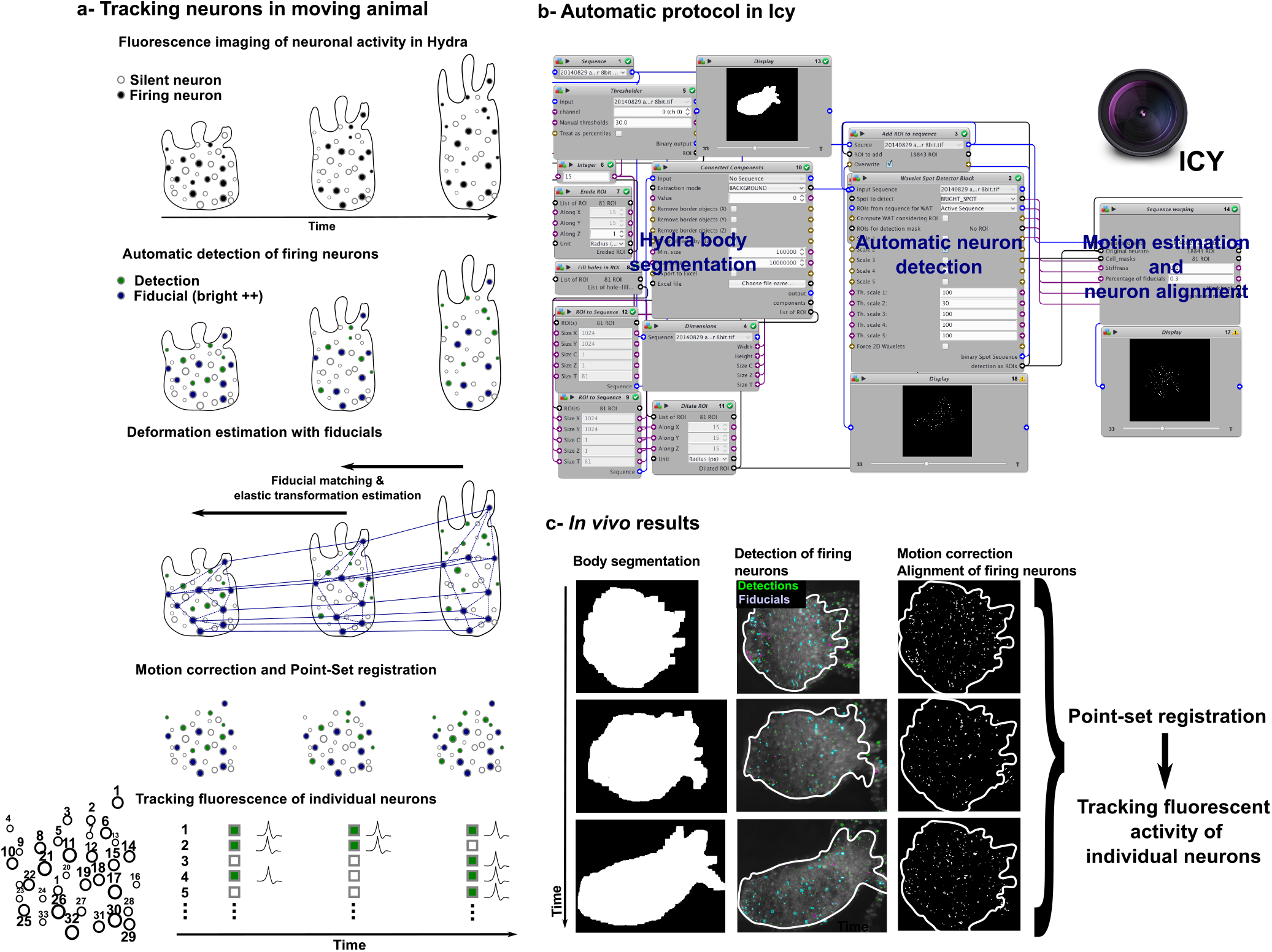
A multi-step algorithm to track individual neuron activity in a deformable animal **a -** General framework of tracking. Neuronal activity is recorded with calcium fluorescent probes. Neurons are homogeneously distributed within Hydra’s deformable body. Firing neurons (bright) are detected with wavelet transformation and statistical thresholding [9]. A fraction of the brightest neurons are used as reference localizations or fiducials for estimating deformation with Thin-Plate Spline (TPS) interpolation. Inverse, iterative transformation of neuron positions allows their alignment with reference frames. As slight neuron motion persists after inverse TPS transform, a point-set registration (Coherent-Point-Drift (CPD) [8]) is applied to track the fluorescence activity of individual neurons. **b -** We designed a protocol in the open-source bio-image analysis platform Icy [10] (http://bioimageanalysis.org/) to automatically perform the multiple analysis steps (Hydra body segmentation, neuron detection and motion correction). The point-set registration (CPD algorithm) was performed by adapting the Python code (https://github.com/siavashk/pycpd) of [8]. **c-** Multi-step tracking approach was applied to time-lapse recordings of Hydra neuronal activity [5]).

## 2. METHODS

### 2.1. Motion correction

To estimate the elastic deformation of Hydra we first automatically detect active, bright neurons using a wavelet-based algorithm [9], implemented as a plugin *Spot Detector* in the open-source image analysis software package Icy [10] (http://icy.bioimageanalysis.org). Big and bright neurons can be robustly detected and tracked over a large number of frames. We thus use a subset (typically 30%) of the brightest detected neurons as references, or *fiducials*, between consecutive frames. These fiducial points are homogeneously distributed over the Hydra body, and can be used to estimate iteratively the elastic deformation of the whole animal. We use the Thin-Plate Spline (TPS) interpolation method [11] whose robustness has been demonstrated in many biological applications, but emphasize that other spline bases such as B-splines [12] might be considered. To align the firing neurons at each time frame with reference frames, we iteratively correct the time-lapse deformation with the estimated TPS interpolation. However, slight motion of individual neurons persists. This, combined with the neuron density and the infrequency of their fluorescence activity, impairs robust tracking. To allow tracking in this case we thus adapt the point-set registration framework (NeRVE) of [6].

### 2.2. Tracking pipeline in motion-corrected animals

Given the motion corrected detections, the NeRVE registration vector clustering method [6] is used to reconstruct neuron tracks. The NeRVE method applies a point set registration from every frame of the video to every frame in a set of reference frames.

After motion correction, a set of spots is extracted from each frame, numbered 1 through N where N varies from frame to frame. The goal of the NeRVE method is then to identify each spot in a given frame with a neuron. For this, one extracts the coordinate of the centroid of each spot for each motion-corrected frame; and these coordinates are collected across many times. Thus, each spot’s identity under each registration to a reference frame is vectorized, and all detected spots in all frames thus acquire a registration vector. To form tracks, the method relies on the intuition that the same neuron in different frames should map in a similar fashion to the set of reference frames. Thus clustering the registration vectors should identify clusters that represent individual neurons. Neuron profiles are then defined as the centers of registration vector clusters, and each spot in each frame is assigned to the neuron with the least Euclidean distance to its registration vector to form the neuron tracks.

After the NeRVE method returns the tracks, we use a post-processing method to fill in gaps by sequentially forming a linear prediction at each frame in the gap, finding the closest detected spot to the prediction, and assigning to the neuron track either the closest spot or the linear prediction if the closest spot falls over a distance threshold. This gives a model-free method for interpolation that accounts for non-linearities in long gaps. Finally, because the accuracy of any point set registration method diminishes as the time between the model (moving) frame and reference (fixed) frames increases, we break long videos into overlapping segments, where the NeRVE method obtains neuron tracks over each segment. Each segment is then spliced together with the subsequent segment by pairing tracks with the least Euclidean error on the overlap.

The point set registration method used in our NeRVE implementation is Coherent Point Drift (CPD) [8]. CPD represents each model point as the centroid of a Gaussian mixture model, and fits the model frame onto the reference frame by maximizing the likelihood of observing the reference points, accounting for a smoothing prior. This posterior is maximized by choosing a set of model points **Y** that minimize the energy function 
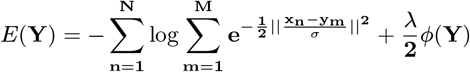

where **X** is the set of reference points, and the *ϕ* function regularizes smoothness of motion. The registration method used in the original NeRVE implementation is a similar Gaussian mixture model registration method [7] which accounts for neuron size and intensity. We chose to use CPD over the Gaussian mixture method because CPD performs better across shorter time frames and all videos can be broken up into short segments using the track splicing method. We also chose CPD for its relative simplicity; the GMM method requires parameters to weigh the relative effects of size and intensity in the energy minimization, and optimal performance with the GMM method also requires a model for how size and intensity change over time. The original NeRVE implementation represents each spot’s image under a registration as a binary vector where an index of 1 represents the closest match under the mapping, with a cut-off parameter that determines when a point has no match under the mapping. As many different representations can be used to capture the global profile of the cell, we chose to use the likelihood vectors (*soft*) given by CPD rather than a binary (*hard*) representation, as it allows us to capture uncertainty in registration and avoid the threshold parameter for deciding that a spot doesn’t have a match.

## 3. COMPARISON WITH MANUAL TRACKING

In order to measure the performance of the tracking method, a ground truth set of tracks was generated in semi-automated fashion. On a 100 frame evaluation video, TrackMate [13] was used to create an initial set of tracks. These were then curated by hand to remove errors and add additional tracks as necessary. The tracks to be evaluated are then associated to the ground truth tracks, allowing for performance statistics to be computed. In order to associate two sets of tracks, we require a way of measuring how *close* they are. We employ the metric used in [14]. Briefly, the metric measures the sum of the Euclidean difference between points, each measured up to a threshold distance *∊*, over all frames. Each frame in which one track is defined but the other is not acquires a penalty of *∊*. Each frame in which neither track is defined acquires no penalty. Thus, for two tracks *θ*^1^ and *θ*^2^,

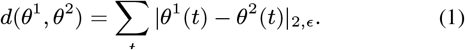

A threshold distance of ∊ = 10 was used. Once a pairwise measure of track distance was defined, the Munkres algorithm [15] was used to find the optimal association between estimated and ground truth tracks. The following per track statistics were computed: (i) Lifetime: length of each associated track, (ii) RMSE: root mean square error over times when both tracks are defined (iii) Proportion less than: proportion of times at which tracks are less than 10 pixels, over times when both tracks are defined. Over all tracks, we also compute the proportion of ground truth tracks that are associated with an estimated track, and the proportion of estimated tracks that are associated with a ground truth track (Fig. 2).

The best results (≈350 over 620 with > 80% detection match) were obtained with motion correction, soft representation of neuron vectors and sequence splicing. We observe that elastic motion correction drastically improves tracking performance. Tracking performances might appear low but, while the ground truth set of tracks are defined at each frame even when neurons are not firing, the automatic tracking depends on bright neuron detections. This leads to incomplete tracks, thus measured statistics should be viewed as comparative rather than absolute performance metrics.

**Fig. 2.**
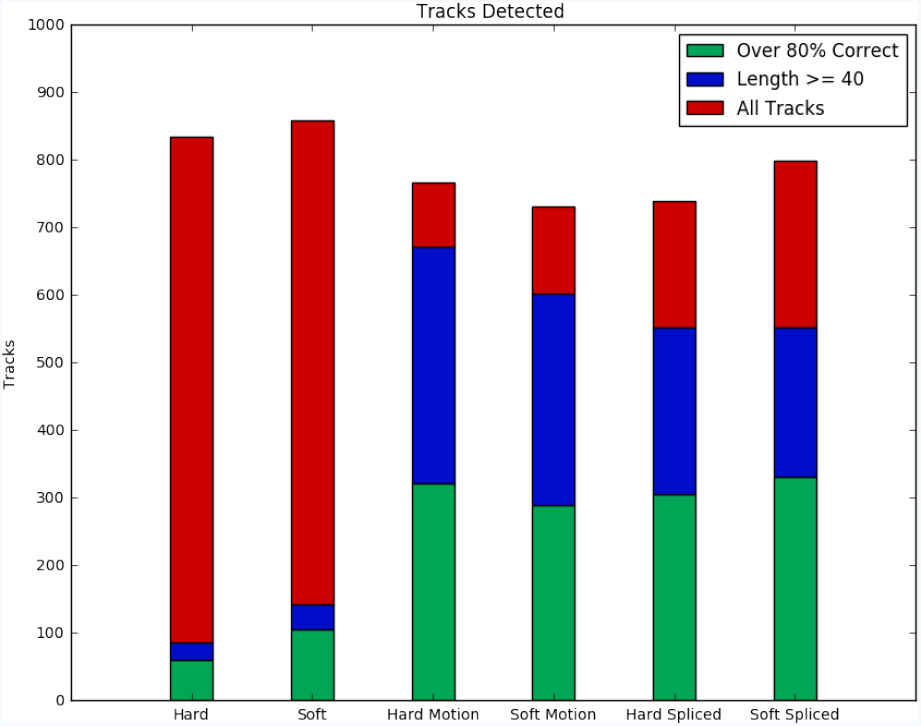
Comparing different tracking strategies with manual tracks (≈ 620) defined over 100 frames. The different compared tracking strategies are, from left to right, (1) hard binary vector representation, without motion correction and sequence splicing, (2) soft probabilistic representation, without motion correction and sequence splicing, (3) hard binary vector representation, with motion correction and without sequence splicing, (4) soft probabilistic vector representation, with motion correction and without sequence splicing, (5) hard binary vector representation, with motion correction and sequence splicing, (6) soft probabilistic vector representation, with motion correction and sequence splicing. Green tracks are the tracks whose length is *>* 40 frames (blue), and more than 80% of the detections match the detections of a ground-truth, manual track. Other short tracks are represented in red.

## 4. CONCLUSION

We proposed a general framework to track neuron activity in the (highly) deformable Hydra. Our strategy consists of (i) identifying very bright neurons that can be unambiguously tracked between consecutive frames, and used as fiducials for estimating the substrate elastic deformation, (ii) iteratively correcting the deformation and aligning firing neurons with reference frames and (iii) using a point-set registration algorithm to robustly track the activity of individual neurons. We compared different tracking strategies with manual tracking (ground truth).

The main contribution of this paper is to propose a general tracking strategy in deformable animals consisting of correcting for deformation with a subset of detected bright neurons (fiducials), before tracking the activity of the entire neuronal population with point-set registration. Each of these steps can be adapted and improved depending on the experimental data-set and the tradeoff with computation load. In the specific example of Hydra, we anticipate additional benefit to come from performing motion correction not just on detected spots but using the entire image frames, a problem that can be solved using a combination of image registration and multi-frame optic flow methods [16, 17]. This allows additional Hydra pose information to be incorporated in motion correction, particularly in cases where the set of active neurons varies in time, and further permits the reliable identification of time points when the Hydra is in a similar position; these identifications should facilitate longer term tracking. This extension is the subject of ongoing work.

## REFERENCES

[1] A. P. Alivisatos, M. Chun, G. M. Church, R. J. Greenspan, M. L. Roukes, and R. Yuste, “A national network of neurotechnology centers for the brain initiative,” Neuron, vol. 88, no. 3, pp. 445–8, Nov 2015.

[2] M. B. Ahrens, J. M. Li, M. B. Orger, D. N. Robson, A. F. Schier, F. Engert, and R. Portugues, “Brain-wide neuronal dynamics during motor adaptation in zebrafish,” Nature, vol. 485, no. 7399, pp. 471–7, May 2012.

[3] R. K. Chhetri, F. Amat, Y. Wan, B. Höckendorf, W. C. Lemon, and P. J. Keller, “Whole-animal functional and developmental imaging with isotropic spatial resolution,” Nat Methods, vol. 12, no. 12, pp. 1171–8, Dec 2015.

[4] J. P. Nguyen, F. B. Shipley, A. N. Linder, G. S. Plummer, M. Liu, S. U. Setru, J. W. Shaevitz, and A. M. Leifer, “Whole-brain calcium imaging with cellular resolution in freely behaving caenorhabditis elegans,” Proc Natl Acad Sci U S A, vol. 113, no. 8, pp. E1074–81, Feb 2016.

[5] C. Dupre and R. Yuste, “Non-overlapping neural networks in hydra vulgaris,” Curr Biol, vol. 27, no. 8, pp. 1085–1097, Apr 2017.

[6] J. P. Nguyen, A. N. Linder, G. S. Plummer, J. W. Shaevitz, and A. M. Leifer, “Automatically tracking neurons in a moving and deforming brain,” PLoS Comput Biol, vol. 13, no. 5, p. e1005517, May 2017.

[7] B. Jian and B. C. Vemuri, “A robust algorithm for point set registration using mixture of gaussians,” in Computer Vision, 2005. ICCV 2005. Tenth IEEE International Conference on, vol. 2. IEEE, 2005, pp. 1246–1251.

[8] A. Myronenko, X. Song, and M. A. Carreira-Perpinán, “Non-rigid point set registration: Coherent point drift,” in Advances in Neural Information Processing Systems, 2007, pp. 1009–1016.

[9] J. C. Olivo-Marin, “Extraction of spots in biological images using multiscale products,” Pattern Recognition, vol. 35, no. 9, pp. 1989–1996, 2002.

[10] F. de Chaumont, S. Dallongeville, N. Chenouard, N. Hervé, S. Pop, T. Provoost, V. Meas-Yedid, P. Pankajakshan, T. Lecomte, Y. Le Montagner, T. Lagache, A. Dufour, and J.-C. Olivo-Marin, “Icy: an open bioimage informatics platform for extended reproducible research,” Nat Methods, vol. 9, no. 7, pp. 690–6, Jul 2012.

[11] J. Duchon, “Splines minimizing rotation-invariant semi-norms in sobolev spaces,” Constructive theory of functions of several variables, pp. 85–100, 1977.

[12] C. O. S. Sorzano, P. Thévenaz, and M. Unser, “Elastic registration of biological images using vector-spline regularization,” IEEE Transactions on Biomedical Engineering, vol. 52, no. 4, pp. 652–663, 2005.

[13] J.-Y. Tinevez, N. Perry, J. Schindelin, G. M. Hoopes, G. D. Reynolds, E. Laplantine, S. Y. Bednarek, S. L. Shorte, and K. W. Eliceiri, “Trackmate: An open and extensible platform for single-particle tracking,” Methods, vol. 115, pp. 80–90, Feb 2017.

[14] N. Chenouard, I. Smal, F. de Chaumont, M. Maška, I. F. Sbalzarini, Y. Gong, J. Cardinale, C. Carthel, S. Coraluppi, M. Winter, A. R. Cohen, W. J. Godinez, K. Rohr, Y. Kalaidzidis, L. Liang, J. Duncan, H. Shen, Y. Xu, K. E. G. Magnusson, J. Jaldén, H. M. Blau, P. Paul-Gilloteaux, P. Roudot, C. Kervrann, F. Waharte, J.-Y. Tinevez, S. L. Shorte, J. Willemse, K. Celler, G. P. van Wezel, H.-W. Dan, Y.-S. Tsai, C. Ortiz de Solórzano, J.-C. Olivo-Marin, and E. Meijering, “Objective comparison of particle tracking methods,” Nat Methods, vol. 11, no. 3, pp. 281–9, Mar 2014.

[15] H. W. Kuhn, “The hungarian method for the assignment problem,” Naval Research Logistics Quarterly, vol. 2, no. 1-2, pp. 83–97, 1955. [Online]. Available: http://dx.doi.org/10.1002/nav.3800020109

[16] R. Garg, A. Roussos, and L. Agapito, “A variational approach to video registration with subspace constraints,” International journal of computer vision, vol. 104, no. 3, pp. 286–314, 2013.

[17] J. Revaud, P. Weinzaepfel, Z. Harchaoui, and C. Schmid, “Deepmatching: Hierarchical deformable dense matching,” International Journal of Computer Vision, vol. 120, no. 3, pp. 300–323, 2016.

